# Evaluation of Alginate-Based Bioinks for 3D Bioprinting, Mesenchymal Stromal Cell Osteogenesis, and Application for Patient-Specific Bone Grafts

**DOI:** 10.1101/2020.08.09.242131

**Authors:** Tomas Gonzalez-Fernandez, Alejandro J. Tenorio, Kevin T. Campbell, Eduardo A. Silva, J. Kent Leach

## Abstract

To realize the promise of 3D bioprinting, it is imperative to develop bioinks that possess the biological and rheological characteristics needed for the printing of cell-laden tissue grafts. Alginate is widely used as a bioink because its rheological properties can be modified through pre-crosslinking or the addition of thickening agents to increase printing resolution. However, modification of alginate’s physicochemical characteristics using common crosslinking agents can affect its cytocompatibility. Therefore, we evaluated the printability, physicochemical properties, and osteogenic potential of four common alginate bioinks: alginate-CaCl_2_ (alg-CaCl_2_), alginate-CaSO_4_ (alg-CaSO_4_), alginate-gelatin (alg-gel) and alginate-nanocellulose (alg-ncel) for the 3D bioprinting of patient-specific osteogenic grafts. While all bioinks possessed similar viscosity, printing fidelity was lower in the pre-crosslinked bioinks. When used to print geometrically defined constructs, alg-CaSO_4_ and alg-ncel exhibited higher mechanical properties and lower mesh size than those printed with alg-CaCl_2_ or alg-gel. The physical properties of these constructs affected the biological performance of encapsulated bone marrow-derived mesenchymal stromal cells (MSCs). Cell-laden constructs printed using alg-CaSO_4_ and alg-ncel exhibited greater cell apoptosis and contained fewer living cells 7 days post-printing. In addition, effective cell-matrix interactions were only observed in alg-CaCl_2_ printed constructs. When cultured in osteogenic media, MSCs in alg-CaCl_2_ constructs exhibited increased osteogenic differentiation compared to the other three bioinks. This bioink was then used to 3D print anatomically accurate cell-laden scaphoid bones that were capable of partial mineralization after 14 days of *in vitro* culture. These results highlight the importance of bioink properties to modulate cell behavior and the biofabrication of clinically relevant bone tissues.

## 1. Introduction

Bone autografts and allografts serve as the current gold standard for the repair of extensive bone damage and delayed unions. However, these procedures are limited by the amount of existing viable bone tissue that can be harvested from a patient, immune rejection, tissue morbidity, and lasting pain at the donor site [1]. Tissue engineering strategies combining biomaterials and progenitor cells such as mesenchymal stromal cells (MSCs) are an exciting alternative to bone grafting for the repair of damaged musculoskeletal tissues [2]. MSCs are a promising cell source due to their proliferative capacity, lack of immunogenicity and multilineage differentiation potential. However, the clinical translation of tissue engineered bone grafts has been modest [3]. More recently, additive manufacturing and 3D bioprinting have emerged as transformative technologies that can speed the clinical translation of tissue engineered products with the promise of generating functional, patient-specific organs that resemble the complex architecture of human tissues [4, 5].

Bioinks, defined as cell-laden hydrogels with the necessary physicochemical properties required for biofabrication, are essential for extrusion-based 3D bioprinting [5, 6]. Micro-extrusion is widely employed for cell printing due to its versatility, enabling the printing of a broad range of bioink viscosities and the potential to 3D print clinically relevant constructs within a realistic time frame [5, 7]. However, the development of hydrogels that provide the required fidelity for high printing resolution while maintaining cell viability, proliferation, and differentiation represents a major limitation for the progress of this technology [5, 6]. A wide range of natural and synthetic hydrogel bioinks with suitable rheological properties have been developed. These bioinks usually contain high polymer concentrations or crosslink densities that increase their viscosity, exhibit shear-thinning behavior, preserve filament shape after extrusion, and possess increased stiffness [5, 8]. However, maximal printing fidelity and high material density may not be optimal to support the cell functions required in tissue engineering [9, 10].

Alginate is an FDA-approved natural polysaccharide that is widely used as a cell carrier, scaffold, and bioink due to its tailorable degradation kinetics, ease of gelation, and possible functionalization with cell adhesive ligands [4, 11]. Prior to crosslinking, alginate solutions behave as non-Newtonian fluids with low viscosities that are unable to acquire a 3D geometrically defined structure without containment inside a mold. Therefore, effective 3D printing of alginate requires that it is modified through crosslinking or by addition of thickening agents to facilitate extrusion as a filament capable of retaining its form. The earliest reports of alginate used for 3D bioprinting focused on the extrusion of a cell-laden alginate solution pre-crosslinked with calcium chloride (CaCl_2_), which was extruded into a CaCl_2_ bath for further crosslinking [12]. Since then, the pre-crosslinking of alginate prior to extrusion with various divalent cations such as CaCl_2_ [13, 14], CaSO_4_ [15], CaCO_3_ [15], BaCO_3_ [16], BaCl_2_ [17] and ZnCl_2_ [17] has been widely used for bone tissue engineering. Despite the simplicity of this approach, poor printing resolution and construct heterogeneity resulting from insufficient control over gelation time, as well as the cellular toxicity of crosslinking agents, have fostered the development of novel hybrid multicomponent bioinks in which pre-crosslinking is no longer needed. The combination of alginate with other natural and synthetic materials such as cellulose [18-20], gelatin [9, 21, 22], hyaluronic acid [23] and Pluronic F-127 [24], among other thickening agents, has resulted in bioinks with optimal viscosity and high cell viability post-printing.

While the characterization of these bioinks is usually reduced to their rheological properties and short-term cell compatibility, a comparative analysis of the long-term cell viability and potential of these bioinks to support osteogenic differentiation of osteoprogenitor cells is lacking. Therefore, we aimed for the first time to assess and systematically compare the ability of four alginate-based bioinks of similar viscosities (alginate-CaCl_2_, alginate-CaSO_4_, alginate-gelatin and alginate-nanocellulose) to be 3D printed, form geometrically defined constructs, maintain cell viability, and support the osteogenic differentiation of MSCs for the biofabrication of patient-specific bone grafts **(figure 1)**.

**Figure 1.**
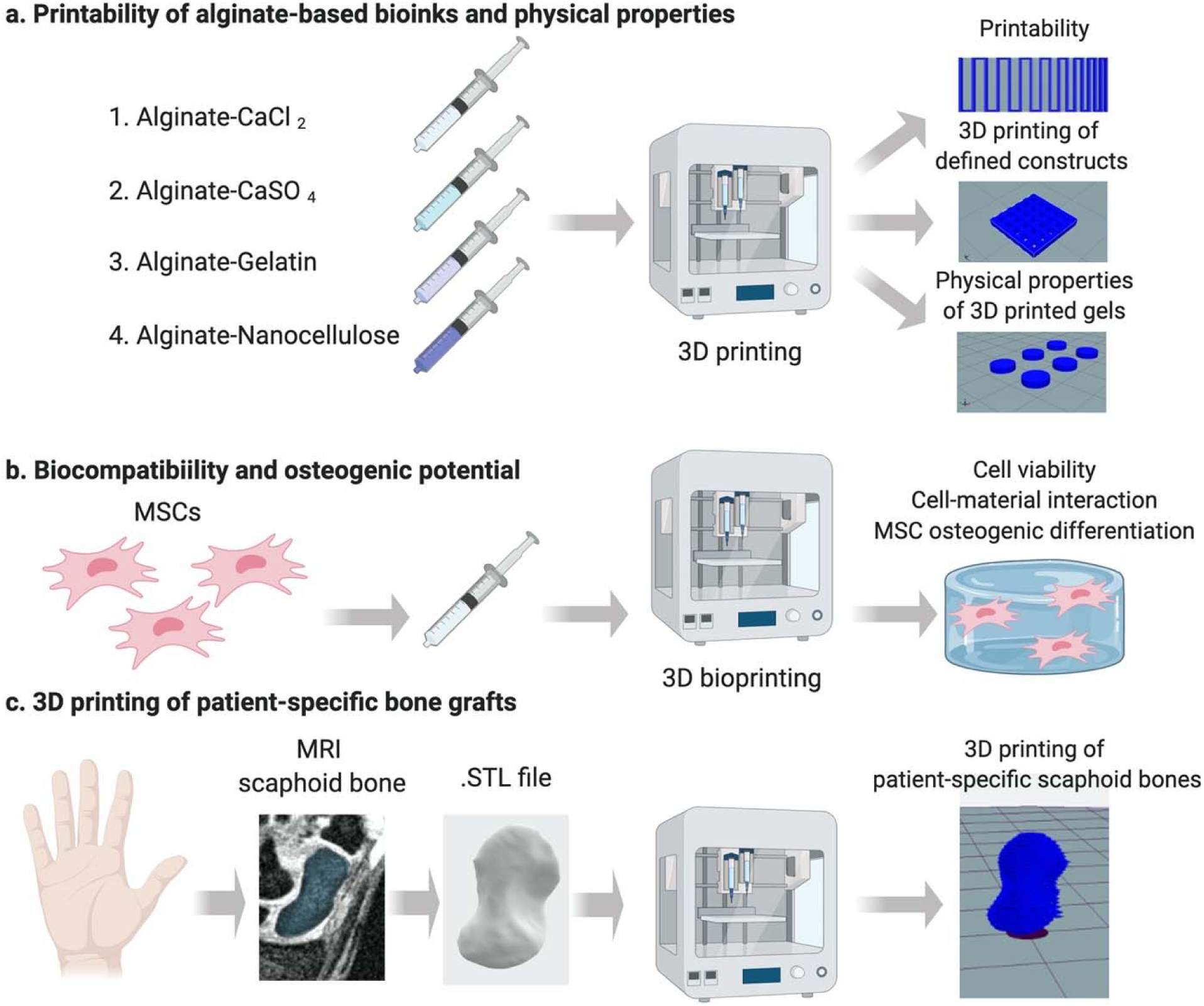
Schematic of experimental plan. **(a)** We compared the printability of 4 alginate-based bioinks (alginate-CaCl_2_, alginate-CaSO_4_, alginate-gelatin, and alginate-nanocellulose). We assessed their potential to form defined constructs and interrogated the physical properties of these constructs. **(b)** The biocompatibility and osteogenic potential of the different bioinks was tested by encapsulating MSCs and evaluating the osteogenic response of cells within 3D-printed constructs. **(c)** Finally, a 3D reconstruction from MRI scans of the scaphoid bone of a healthy patient was bioprinted, and we assessed its potential for the fabrication of cell-laden patient-specific osteogenic grafts.

## 2. Materials and methods

### 2.1. Bioink preparation

Ultrapure MVG sodium alginate (viscosity > 200 mPa.s, MW > 200 kDa and G/M ratio ≥ 1.5; Pronova FMC BioPolymer, Norway) was covalently modified with RGD peptide (GGGGRGDSP; Peptide 2.0, Chantilly, VA) using carbodiimide chemistry as previously reported [25, 26]. The molar ratio of RGD to alginate was varied such that each alginate chain possessed a degree of substitution (DS) of 2. The modified alginate was then lyophilized for one week and resuspended in phosphate-buffered saline (PBS) to a final concentration of 3.5%. For the alginate-CaCl_2_ (alg-CaCl_2_) bioink, CaCl_2_ (MilliporeSigma, Burlington, MA) was prepared in ultrapure water (UPW) at 50 mM and then mixed for 2 min at a 3:7 ratio with the 3.5% alginate solution to obtain a homogeneous mixture. For the alginate-calcium sulfate (alg-CaSO_4_) bioink, calcium sulfate (CaSO_4_, MiliporeSigma) was prepared in UPW at 50 mM and then mixed for 2 min at a 3:7 with the 3.5% alginate solution to obtain a homogeneous mixture. For the alginate-gelatin (alg-gel) bioink, gelatin (type B from bovine skin, MilliporeSigma) was dissolved in PBS for 10 min at 37°C to form a 15% solution, which was then mixed for 2 min at a 3:7 ratio with the 3.5% alginate solution to obtain a homogeneous mixture and incubated at 4°C for 10 min to allow the gelatin to crosslink. Alginate-nanocellulose (alg-ncel) bioink was purchased from CELLINK (IK1020000301, Boston, MA).

### 2.2. Bioink rheological characterization

The rheological properties of each bioink were characterized using a Discovery HR-2 hybrid stress-controlled rheometer (Thermal Analysis Instruments, New Castle, DE) equipped with a 40 mm parallel plate geometry and a measurement gap of 0.55 mm. Shear rates in the range of 0.1 to 1000 s^-1^ at a frequency of 1 Hz were used in order to determine the linear viscoelastic range of the bioinks. To assess the recovery of the viscosity of the materials and their thixotropic behavior, a recovery test was performed by subjecting the bioinks to 1 s^-1^ shear rate for 60 s, 100 s^-1^ shear rate for 10 s and 1 s^-1^ shear rate for 60 s to simulate the printing process [27], where a shear rate of 100 s^-1^ is estimated as the maximum shear rate experienced by the hydrogel during printing [27]. 2.5% alginate solution and Pluronic F127 (Pluronic, MilliporeSigma) were used as negative and positive controls, respectively.

### 2.3. Bioink 3D printing and printability characterization

Bioinks were printed using an Allevi 2 3D bioprinter (Allevi, Philadelphia, PA). Bioinks were loaded into the pneumatic driven syringes equipped with 25 Gauge (G) and 0.25’’ needles (Allevi). To assess printing fidelity, the filament fusion test was performed at pressures of 10, 20 and 30 psi and followed a printing pattern with a filament distance of 0.25 mm, increased 0.05 mm for each subsequent line, and finishing at the distance of 1.0 mm as previously described [28] **(figure 2(c))**. Printability was optimized for each bioink by identifying the pressure which resulted in the highest resolution and smallest filament spreading ratio, defined as the width of the printed filament divided by the needle diameter [29]. The filament collapse test was performed using a 3D printed platform as described [28] in which pillars (l × w × h = 2.0 × 2.0 × 4.0 mm) were placed at known gap distances (1.0, 2.0, 4.0, 8.0 and 16.0 mm) between each other **(figure 2 (d))**. The spreading ratio and the filament collapse test were analyzed using ImageJ software.

**Figure 2.**
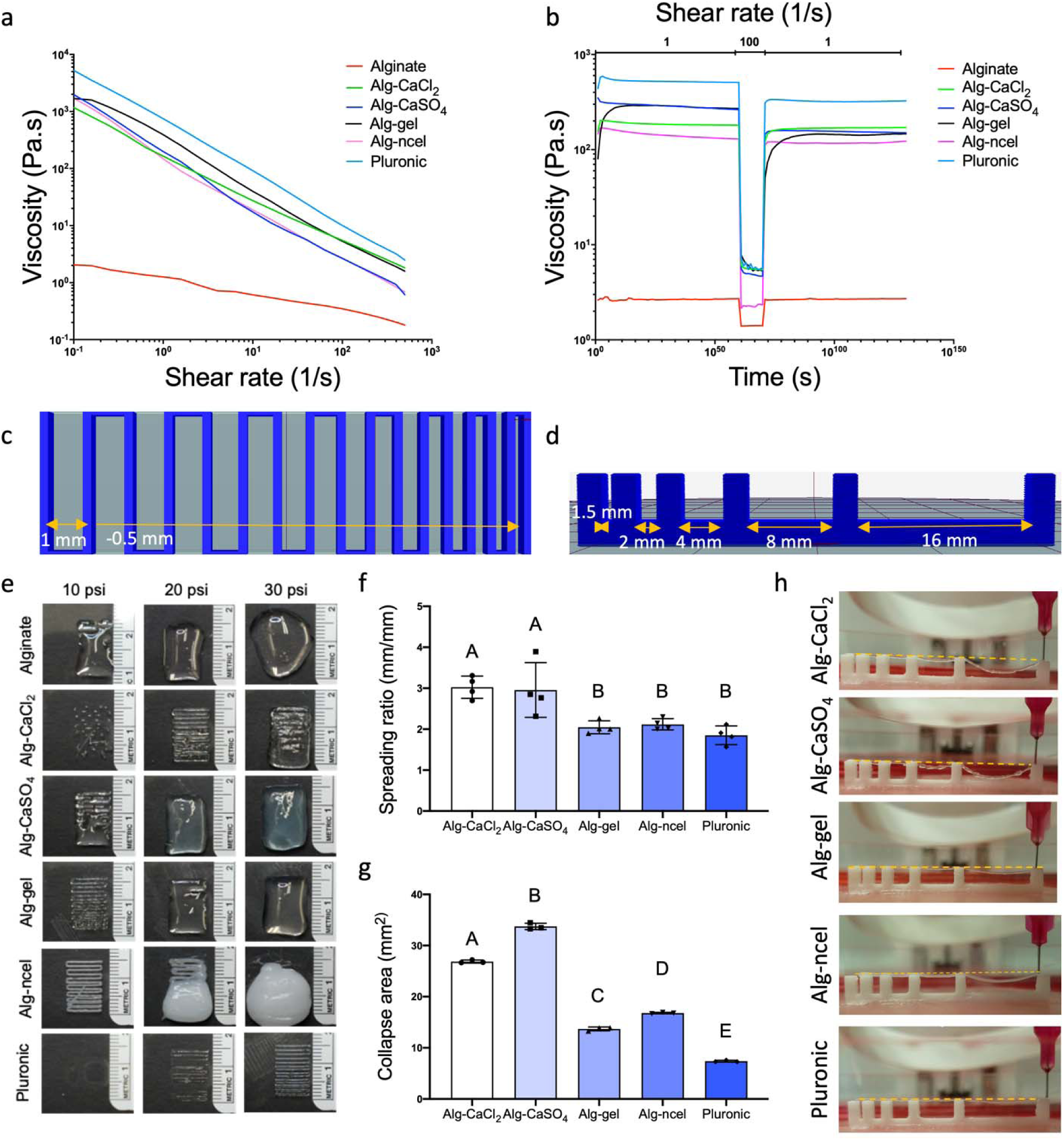
Rheological properties and printability of alginate-based bioinks. **(a)** Viscosity and **(b)** thixotropic behavior of the four bioinks. Schematics of the **(c)** filament fusion printing pattern and **(d)** filament collapse platform used to determine bioink printability. **(e)** Images of the filament fusion test of the four bioinks printed at 10, 20 and 30 psi. Quantification of **(f)** filament spreading ratio (n = 4) and **(g)** collapse area of the bioinks (n = 3). **(h)** Filament collapse images of the different bioinks.

### 2.4. Characterization of mechanical and physical properties of 3D printed constructs

Macroporous square constructs **(figure 1(a))** of 2 cm × 2 cm × 0.5 cm and 1 cm × 1cm × 0.25 cm and with an internal structure of 25 square pores of 2 mm × 2 mm and 0.1 cm × 0.1 cm, respectively, were 3D printed and crosslinked with 100 mM CaCl_2_ for 5 min. Constructs were imaged before and after final crosslinking. To assess the change in physical and mechanical properties over time, constructs with 8 mm diameter x 2 mm height **(figure 1(a))** were 3D printed, crosslinked in sterile conditions, and kept in growth media consisting of minimum essential alpha medium (αMEM; Invitrogen, Carlsbad, CA) supplemented with 10% fetal bovine serum (FBS; Atlanta Biologicals, Flowery Branch, GA) and 1% penicillin/streptomycin (Gemini Bio Products, Sacramento, CA) at 37°C. At t = 0 and 7 days, constructs were removed to determine storage modulus, swelling ratio, and mesh size. To assess the storage modulus of 3D printed constructs, constructs were tested using a Discovery HR-2 hybrid stress-controlled rheometer. The constructs were placed between 8 mm parallel plates (axial force set at 0.2 N) and were strained from a range of 0.001-5% with a frequency of 1 Hz. The values of storage modulus (*G*′) were obtained from the linear viscoelastic region. In order to obtain the volumetric swelling ratio, the wet weight (*w*_*s*_) of the constructs were recorded, the constructs were frozen at -80°C overnight, lyophilized for 48 h, and then weighed again to obtain the dry weight (*w*_*D*_). Volumetric swelling ratio (*Q*) was calculated by the following equation as previously described [30, 31]:

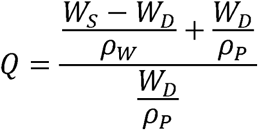

where ρ_*W*_ is the density of water and ρ_*p*_ is the density of polymer (1.6 g/mL). Mesh size of the crosslinked polymers in the constructs was estimated using swelling and storage modulus as previously described [30-32]. The molecular weight between crosslinks (*M*_C_) was then determined *via* the following equation:

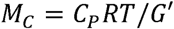

where *C*_*p*_ is the polymer concentration, *R* is the gas constant and *T* is temperature. Mesh size (ξ) was then calculated as follows:

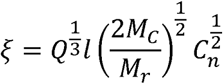

Where *l* is the length of the repeating unit (5.15 Å), *M*_*r*_ is the molecular weight of the repeating unit (194 g/mol) and *c*_*n*_ is the characteristic ratio (21.1).

### 2.5. Cell culture and 3D bioprinting of cell laden bioinks

Human bone marrow-derived mesenchymal stromal cells (MSCs) from a single donor were purchased from Lonza (Walkersville, MD). The trilineage potential of the cells was confirmed through induction in lineage-specific media. MSCs were expanded under standard conditions at 37°C and 21% O_2_ in growth medium until use at passage 4. Media was refreshed every 2-3 days. After reaching 80% confluency, MSCs were trypsinized with Trypsin-EDTA (0.25%) (Thermo Fisher, Waltham, MA) and counted using trypan blue exclusion staining with a Countess II automatic cell counter (Thermo Fisher). 10×10^6^ cells/mL were encapsulated in each bioink and thoroughly mixed for 2 min. For the alg-gel bioink only, the bioink was incubated at room temperature for 30-45 min after mixing to enable gelatin crosslinking. Cell-laden bioinks were used to 3D print cylindrical constructs of 4 mm diameter x 1.5 mm height. Immediately after printing, cell-laden constructs were crosslinked for 5 min in 100 mM CaCl_2_ dissolved in growth media, after which constructs were transferred to standard growth media.

For osteogenic induction, constructs were incubated in growth media for 24 h after printing and then changed to osteogenic medium composed of growth media supplemented with 50 μM ascorbate 2-phosphate, 10 mM β-glycerophosphate, and 100 nM dexamethasone (all from Sigma-Aldrich, St. Louis, MO) for 14 days under standard conditions. Media was changed every 2-3 days.

### 2.6. Assessment of cell viability

Cell viability was analyzed by a live/dead assay per the manufacturer’s protocol (Thermo Fisher). Fluorescent images were taken using confocal microscopy (Leica TCS SP8, Wetzlar, Germany) and the percentage of living cells was quantified using ImageJ. ImageJ was also used for the analysis of cell shape descriptors. For quantification of DNA level and caspase 3/7 activity, constructs were collected, sonicated, and lysed in passive lysis buffer (Promega, Madison, WI). Total DNA content was determined using a PicoGreen Quant-iT PicoGreen DNA Assay Kit (Invitrogen). Cell apoptosis was measured using a Caspase-Glo 3/7 assay (Promega, Madison, WI).

### 2.7. Assessment of cell morphology

Constructs were collected after 7 days of culture and fixed with 4% paraformaldehyde at 4°C overnight, washed twice with PBS, and permeabilized with 0.05% Triton-X 100 for 5 min at room temperature. Cell actin cytoskeleton was stained with Alexa Fluor 488 Phalloidin solution (Thermo Fisher; 1:40 in PBS). Cell nuclei were stained with DAPI (Thermo Fisher; 1:500 in PBS). Gels were imaged using confocal microscopy (Leica TCS SP8).

### 2.8. Biochemical and histological evaluation of osteogenic constructs

Calcium content was determined using a Stanbio Calcium Liquid Reagent for Diagnostic Set (Thermo Fisher) after digestion in 1M HCl at 60°C for 72 h. Total collagen content was determined by measuring hydroxyproline content using the dimethylaminobenzaldehyde and chloramine T assay and a hydroxyproline to collagen ratio of 1:7.69 [33].

For histological evaluation of 3D printed constructs, constructs were fixed in 4% PFA overnight at 4°C and washed in UPW. Samples were dehydrated in a graded series of ethanol baths and paraffin-embedded overnight. Each gel was sectioned at 7 μm thickness using a Leica RM2235 Manual Rotary Microtome and affixed to microscope slides for subsequent staining. The sections were stained with H&E to assess cell-matrix interaction, picrosirius red to determine collagen content, and alizarin red to observe calcification. Osteocalcin was detected using a primary antibody against osteocalcin (Abcam, Cambridge, MA; AB13420; 1:200).

### 2.9. 3D printing of cell-laden patient-specific scaphoid bone grafts

3D reconstructions of the scaphoid of a healthy volunteer were provided by Prof. Abhijit J. Chaudhari (UC Davis, Department of Radiology) from MRI scans with a spatial resolution of 0.45 × 0.45 × 0.50 mm. The 3D reconstruction .stl files were sliced with the open source Slic3r software (slic3r.org US) in Repetier-Host to generate G-code files. Alg-CaCl_2_ bioinks were produced as described above, and MSCs were encapsulated at 10×10^6^ cells/mL. Immediately after printing, alg-CaCl_2_-printed scaphoids were further crosslinked with 100 mM CaCl_2_ for 5 min in 100 mM CaCl_2_ dissolved in growth media, after which constructs were transferred to fresh growth media for 24 h and cultured in osteogenic media for 14 days.

### 2.10. Statistical analysis

Data are presented as means ± standard deviation. Statistical analysis utilized a one-way ANOVA with post-hoc Tukey test. *p* < 0.05 was considered significant. In each graph, data points with different letters are significantly different from one another.

## 3. Results and Discussion

### 3.1. Alginate-based bioinks with similar viscosity and thixotropic behavior have different printability

Bioink printability is not unequivocally defined and lacks an accepted, standardized method for its analysis [28, 34]. The printability window is often predicted by bioink rheological properties such as viscosity, shear thinning, and recovery behavior to establish optimal printing parameters for the application [35]. Although the analysis of these properties is critical for understanding the parameters that guide bioink optimization, they are insufficient to determine printability [34, 36]. The characterization of bioink shape fidelity after printing is needed to assess physical deformation of the printed filament and degree of similarity to the CAD design [28]. In this study, the four different alginate-based bioinks possessed overlapping linear viscosities that were higher than 2.5% alginate but lower than Pluronic **(figure 2(a))**, which was included as a positive control due to its excellent printability [37]. Bioink thixotropy, characterized through a viscosity recovery test, revealed greater shear thinning behavior of the bioinks compared to 2.5% alginate **(figure 2(b))**.

We performed the filament fusion test [28] at 10, 20 and 30 psi to assess bioink shape fidelity after extrusion and identify an adequate printing pressure range for each bioink **(figure 2(c) and (e))**. While a higher pressure of 20 psi was needed for the alg-CaCl_2_ bioink, the remaining bioinks were printable at 10 psi **(figure 2(e))**. The increased pressure could be due to the rapid and uncontrolled crosslinking of alginate caused by the quick release of Ca^2+^ ions, forming heterogeneous crosslinks within the bioink [22, 38] and necessitating higher pneumatic pressures for extrusion of a continuous filament. Despite being pre-crosslinked with the same concentration of Ca^2+^, alg-CaSO_4_ was printable at lower extrusion pressures than alg-CaCl_2_, perhaps due to the slower gelation kinetics caused by the lower solubility of sulfate salts [39]. Lower CaSO_4_ solubility could also explain similar results observed by Freeman *et al*., in which higher printing pressures were needed to extrude alg-CaCl_2_ bioinks in comparison to alg-CaSO_4_ and alg-CaCO_3_ [15]. The filament fusion test was also used to calculate the bioink spreading ratio to assess filament deformation [29]. Alg-gel and alg-ncel showed the lowest spreading ratios, similar to Pluronic, while alg-CaCl_2_ and alg-CaSO_4_ exhibited significantly higher ratios, suggesting lower printing fidelity **(figure 2(f))**. The filament collapse test was also performed to assess filament deflection when suspended **(figure 2(d))** [28]. Similar to the spreading ratio, alg-gel and alg-ncel possessed the lowest collapse area of the four bioinks **(figure 2(g) and (h))**. The decreased printability of pre-crosslinked alginate bioinks could be attributed to their heterogeneous crosslinking densities. Chung *et al*. found higher variability in the extrusion force used for alg-CaCl_2_ bioinks than alg-gel bioinks, indicating higher heterogeneity within the alg-CaCl_2_ solution and thus lower printing resolution [22].

### 3.2. Physical properties of 3D printed construct depend on the type of alginate-based bioink

The printability of the four bioinks and capacity to produce tissue grafts with a defined architecture were confirmed through the 3D printing of geometrically defined, macroporous constructs. Bioinks were further crosslinked after printing in a CaCl_2_ bath to maintain the construct shape **(figure 3(a) and (b))**. Macroscopic evaluation revealed that all 3D printed 1cm x 1cm constructs maintained their internal architecture after crosslinking, with the exception of the alg-CaSO_4_ constructs **(figure 3(c) and (d))**. The printing of these constructs also facilitated analysis of the internal filament structure through H&E staining, which confirmed the heterogeneity of the pre-crosslinked bioinks compared to the homogenous structure of the alg-gel and alg-ncel **(figure 3(d))**.

**Figure 3.**
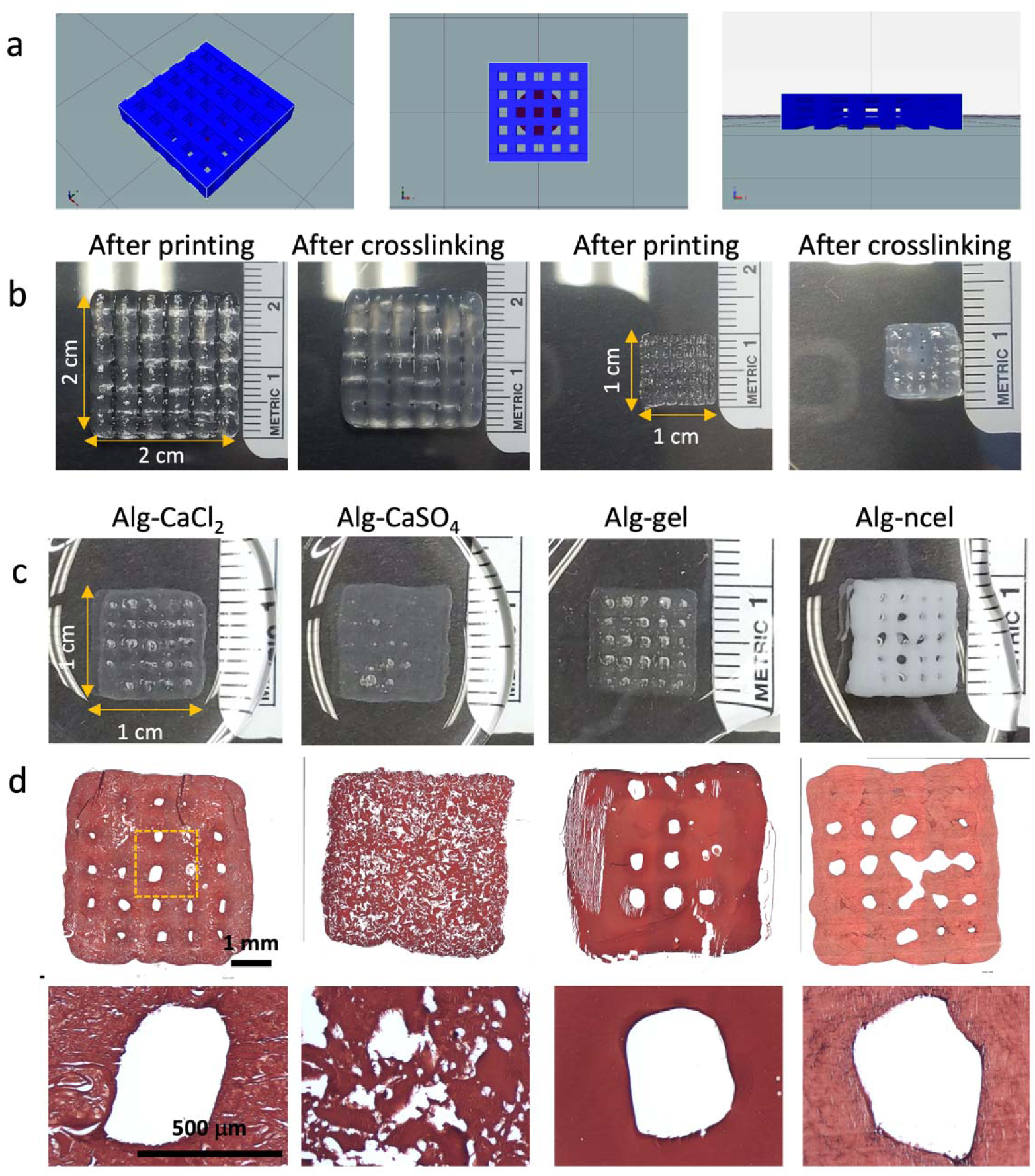
3D printing of geometrically defined macroporous constructs. **(a)** Schematic of geometrically defined macroscopic construct formed from Slic3r software. **(b)** Macroscopic images of 3D printed alg-CaCl_2_ constructs of different sizes (2 cm x 2 cm and 1 cm x 1 cm) with defined geometry and porosity immediately after printing and after crosslinking for 5 min in a CaCl_2_ bath. **(c)** Macroscopic images of 3D printed geometrically defined constructs with the four different alginate-based bioinks after crosslinking in a CaCl_2_ bath. **(d)** Histological examination of construct microscopic structure by H&E staining reveals maintenance of pore structure.

We assessed the physical properties of geometrically defined printed constructs (8 mm diameter x 2 mm height cylindrical constructs) **(figure 4(a) and (b))** after crosslinking in a CaCl_2_ bath. Physical properties were assessed after printing (day 0) and after 7 days of *in vitro* culture (day 7). The mechanical properties of alginate hydrogels are key for controlling cell adhesion and differentiation [40]. In this study, alg-CaSO_4_ and alg-ncel bioinks formed constructs that possessed an initially higher storage modulus than alg-CaCl_2_ and alg-gel constructs (9.9 ± 4.9 kPa for alg-CaSO_4_, 7.1 ± 0.6 kPa for alg-ncel, 4.1 ± 0.6 kPa for alg-CaCl_2_, and 2.6 ± 1.1 kPa for alg-gel). After 7 days in culture, all hydrogels exhibited a decrease in stiffness except the alg-ncel group that retained a consistent modulus **(figure 4(c))**. This decrease in mechanical properties in the alg-CaCl_2_ and alg-CaSO_4_ constructs over 7 days could be the result of the dissociation of calcium crosslinks [41] rather than by the degradation of the alginate polymer, as a significant decrease in dry weight was only observed in the alg-gel groups (*data not shown*). In alg-gel hydrogels, the decrease in dry weight was likely due to gelatin reverting to its soluble form at physiological temperatures, resulting in a loss of the polymer network and a corresponding reduction in storage modulus. The swelling behavior and mesh size also play an important role in regulating nutrient transport, cell-cell and cell-substrate interactions, cell viability, and proliferation [42, 43]. Analysis of the volumetric swelling ratio initially revealed significantly lower swelling for the alg-gel and alg-ncel constructs compared to the alg-CaCl_2_ and alg-CaSO_4_ constructs (34.1 ± 3.1 for alg-CaCl_2_, 36.2 ± 4.1 for alg-CaSO_4_, 25.4 ± 1.7 for alg-gel and 26.4 ± 2.2 for alg-ncel, *p*<0.0001 for alg-CaSO_4_ *vs* alg-gel and alg-ncel, *p* = 0.0003 and *p* = 0.0011 for alg-CaCl_2_ *vs* alg-gel and alg-ncel, respectively), though only the alg-gel constructs had an increase in swelling after 7 days **(figure 4(d))**. Interestingly, mesh size was higher in the alg-CaCl_2_ and alg-gel constructs at day 0 (80.5 ± 8.4 nm for alg-CaCl_2_, 56.7 ± 12.8 nm for alg-CaSO_4_, 96.6 ± 15.6 for alg-gel, and 56.2 ± 2.9 for alg-ncel, *p* = 0.0067 and *p* = 0.0056 for alg-CaCl_2_ *vs* alg-CaSO_4_ and alg-ncel respectively, *p*<0.0001 for alg-gel *vs* alg-CaSO_4_ and alg-ncel). The mesh size increased over 7 days in all gels except alg-ncel constructs, which exhibited a slight decrease in mesh size (**figure 4(e))**. Taken together, these results demonstrate how the selection of bioinks influences the properties of 3D printed constructs.

**Figure 4.**
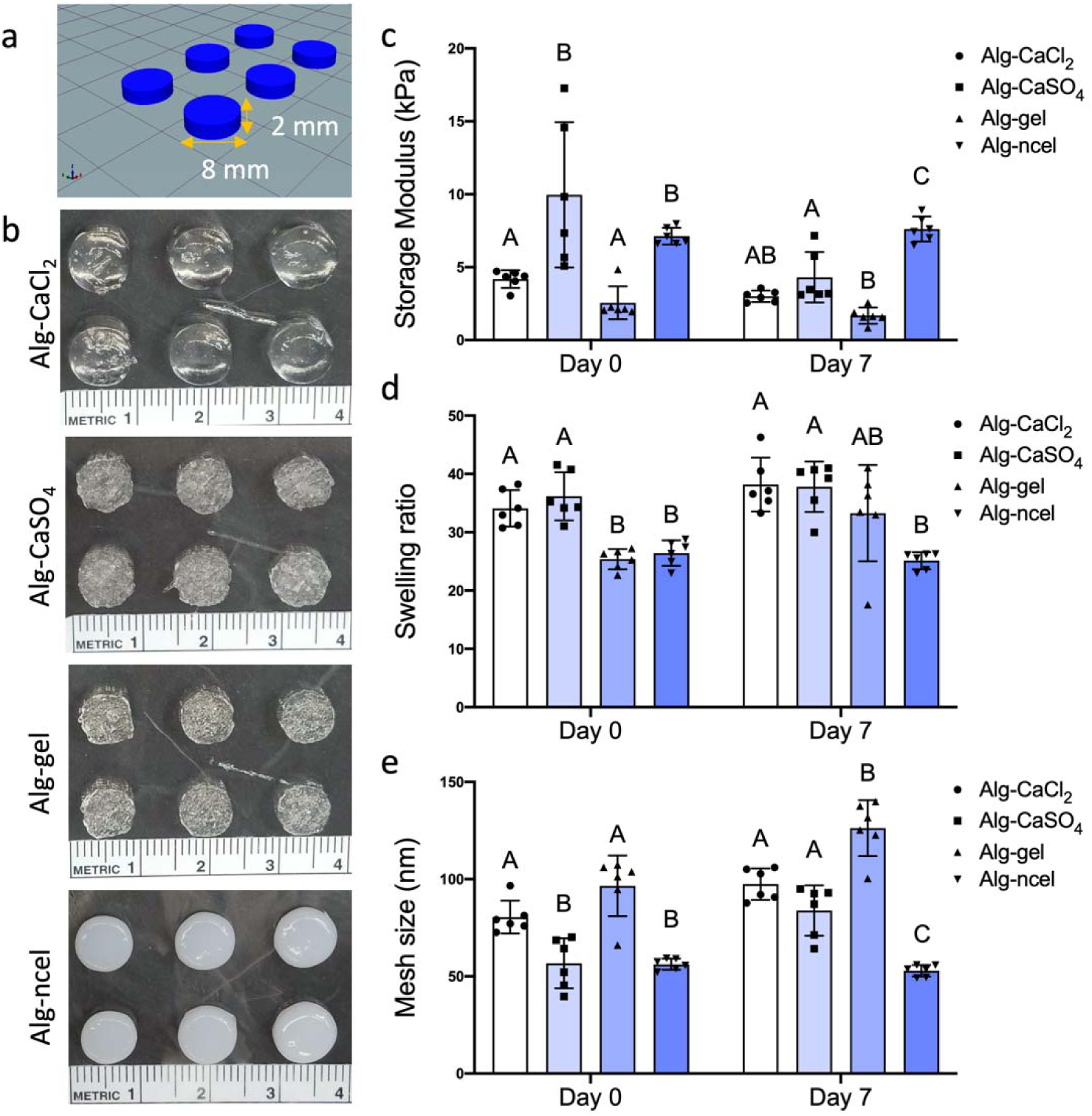
Biophysical properties of geometrically defined 3D printed cylindrical constructs. **(a)** Schematic of 3D printed cylindrical construct. **(b)** Macroscopic images of 3D printed cylindrical constructs with the four different alginate-based bioinks. **(c)** Storage modulus of 3D printed constructs after printing (day 0) and after 7 days in culture (day 7). **(d)** Swelling ratio of the 3D printed cylinders at days 0 and 7. **(e)** Mesh size of the 3D printed constructs at days 0 and 7 (n = 6 for (c), (d) and (e)).

### 3.3. Stromal cell behavior in 3D bioprinted constructs depends on the type of bioink

The 3D printing of living tissues requires the maintenance of cell viability during and after the printing process. While cell viability immediately after printing depends on bioink preparation and the shear stresses generated during the 3D printing process, the long-term cytocompatibility is determined by construct properties. At day 1 after printing cylindrical gels (4 mm diameter by 1.5 mm height), all groups exhibited similar DNA content **(figure 5(a))**, but the percentage of living cells was lower in the alg-CaCl_2_ and alg-gel constructs **(figure 5(c) and (d))**. These differences could be due to the greater extrusion pressure required for the 3D printing of alg-CaCl_2_ bioinks [44] (>20 psi *vs*. 10-15 psi used in the other bioinks). In the case of the alg-gel bioink, differences in cell viability may be due to the incubation time of 30-45 minutes at room temperature after mixing to ensure gelatin crosslinking before printing as previously described [9], while the remaining bioinks were used immediately after mixing. Although the alg-CaCl_2_ and alg-gel constructs showed the lowest percentage of living cells at day 1, cells in these gels had the lowest levels of apoptosis *via* caspase 3/7 activity **(figure 5(b))**. This difference could be due to the low half-lives of caspases 3 and 7 (8 and 11 hours, respectively [45]), suggesting that cell apoptosis 24 hours after printing depends solely on material properties. After 7 days of culture, we observed a significant increase in cell apoptosis and decrease in living cells in alg-CaSO_4_ and alg-ncel constructs **(figure 5(a-c))**, which may be due to the initially higher storage modulus and lower mesh sizes of these hydrogels **(figure 4)**, potentially limiting nutrient transport to encapsulated cells. Although an increase in apoptosis and cell death was not observed in alg-gel constructs, these gels possessed the lowest DNA content **(figure 5(h))** at day 7. Previous work has shown that cells can displace the polymer system of weaker alginate hydrogels [30], suggesting that the MSCs in these constructs potentially displaced the polymer network and migrated from the lower storage modulus and higher mesh size alg-gel 3D printed hydrogels.

**Figure 5.**
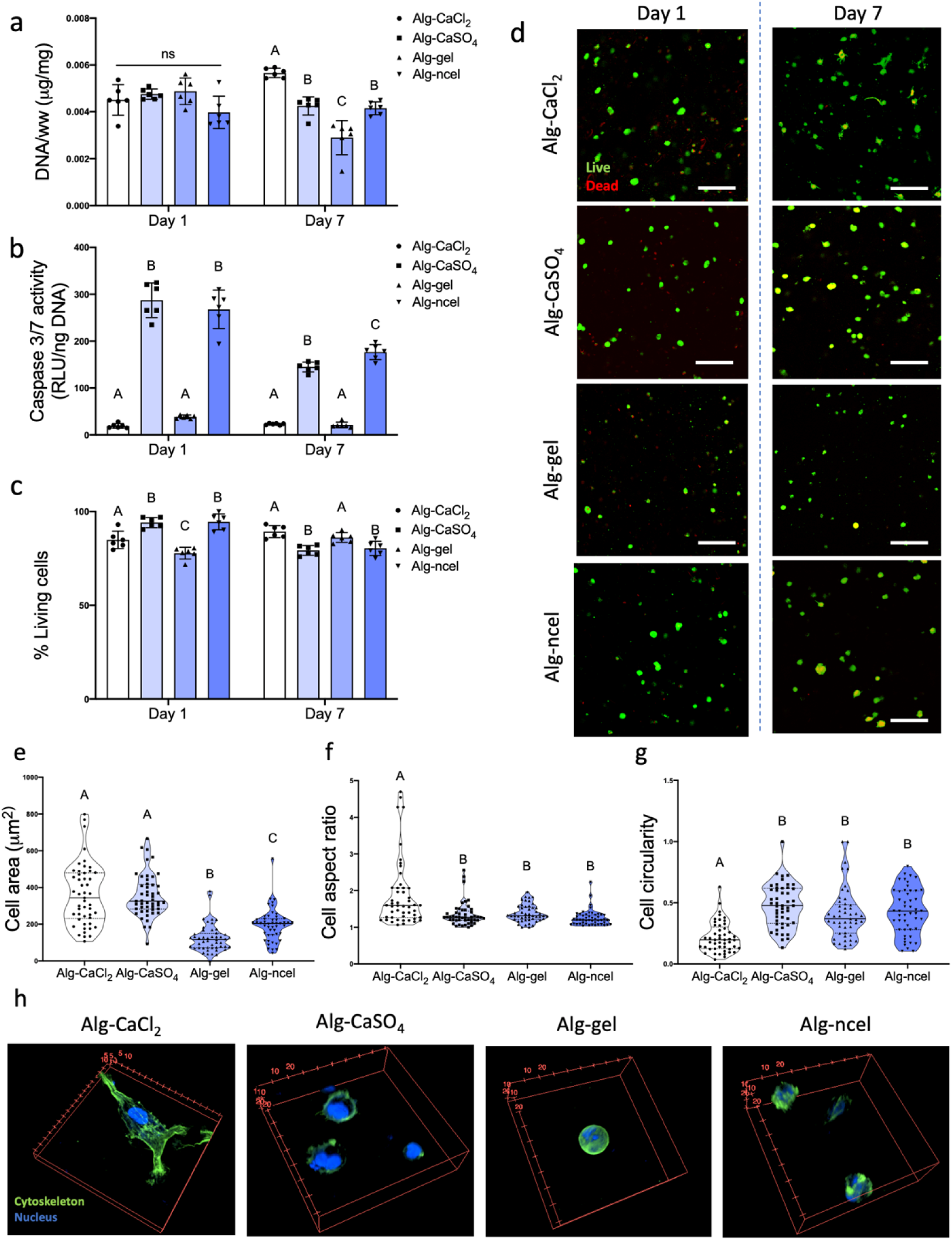
MSC viability, metabolic activity, and cell-substrate interaction depend on the type of alginate-based bioink. **(a)** DNA content in 3D printed constructs at day 1 and 7 after biofabrication. **(b)** Caspase 3/7 activity in the 3D printed constructs at day 1 and 7 after biofabrication. **(c)** Percentage of living cells in the 3D printed gels at day 1 and 7 after printing (n = 6 for panels a-c). **(d)** Live/dead confocal images of cells in 3D printed gels at days 1 and 7 after printing (live cells are green, dead cells are red). Quantification of **(e)** cell area, **(f)** cell aspect ratio, and **(g)** cell circularity as shape descriptors of cells encapsulated in 3D printed cells at day 7 after printing (n = 49 for panels e-g; ImageJ was used to analyze 3 images per group). **(h)** 3D confocal reconstructions of cells encapsulated in 3D printed constructs and stained for their actin cytoskeleton (green) and nucleus (blue). Scale bar = 200 μm.

Alginate lacks endogenous motifs to enable cell adhesion. Modification of the alginate backbone with the fibronectin-derived Arginine-Glycine-Aspartic Acid (RGD) peptide has been widely explored to promote cell adhesion and enhance MSC osteogenesis [4, 46, 47]. Live/dead staining of encapsulated cells revealed differences in cell morphology and size at day 7 after printing **(figure 5(d))**. Quantification of cell size and shape revealed larger cells in the alg-CaCl_2_ and alg-CaSO_4_ printed gels **(figure 5(e))**. The highest cell aspect ratio and lowest circularity, as indicators of cell elongation, were detected in alg-CaCl_2_ constructs **(figure 5(f) and (g))**. To further observe the effects of bioink encapsulation on cell morphology and actin cytoskeleton, 3D printed hydrogels containing MSCs were stained with phalloidin after 7 days of culture **(figure 5(h))**. Fluorescent confocal imaging revealed more elongated cells with an organized cytoskeleton and well-defined actin fibers only in the alg-CaCl_2_ hydrogels, suggesting stronger interaction of the encapsulated cells with the RGD ligand on the alginate polymer **(figure 5(h))**. We observed limited cell spreading in alg-CaSO_4_ constructs compared to alg-CaCl_2_, which may be due to cell confinement due to higher crosslinking density and lower mesh size in the alg-CaSO_4_ constructs **(figure 4(e))** [48]. Similarly, the lack of MSC spreading in the alg-ncel printed gels could be due to low mesh size, as cells encapsulated in alg-ncel bioinks achieve a spread morphology independently of RGD modification [19, 20]. Although alg-gel printed constructs possessed mesh sizes and mechanical properties similar to alg-CaCl_2_ **(figure 4)**, we did not observe cell spreading in this bioink, likely due to the reversion of the gelatin to its soluble form at 37°C and the inability of cells to bind to denatured collagen [49].

### 3.4. MSC osteogenic potential depends on the type of alginate-based bioink

We analyzed the osteogenic potential of MSCs encapsulated in the four alginate-based bioinks. Osteogenesis of MSCs in alginate gels is governed by a combination of substrate mechanical properties and cell-matrix interactions through integrin–adhesion ligand binding. While MSCs entrapped in softer alginate hydrogels (Young’s Modulus of 2.5-5 kPa) undergo adipogenic commitment, MSCs in stiffer gels (11–30□kPa) exhibit increased osteogenic differentiation [40]. The abrogation of RGD binding to α5 integrins has been shown to decrease osteogenesis and enhance adipogenesis in stiff hydrogels, highlighting the role of cell-matrix interaction for MSC differentiation [40]. In this study, despite lower storage modulus **(figure 4(b))**, MSCs encapsulated in 3D printed alg-CaCl_2_ gels achieved the highest calcium deposition **(figure 6(c)** and **(d))** and positive osteocalcin staining **(figure 6(d))**, used as a marker of osteoblastic differentiation, after 14 days of culture. In a previous study, Freeman *et al*., harnessed the higher stiffness of alg-CaSO_4_ 3D printed constructs to promote MSC osteogenesis in comparison to the use of alg-CaCl_2_ as bioink. In contrast, the use of RGD-modified alginate in the current work suggests that early cell adhesion and spreading in the alg-CaCl_2_ hydrogels may have enhanced MSC osteogenesis in comparison to alg-CaSO_4_. Alg-CaSO_4_ and alg-ncel, which possessed similar physical properties **(figure 4)**, also induced similar levels of calcium deposition **(figure 6(c))**. Although MSCs in alg-CaSO_4_ stained positive for osteocalcin, this marker was not observed in the alg-ncel gels **(figure 6(d))**. While cell spreading was not observed in alg-CaSO_4_ constructs at day 7 **(figure 5(h))**, live/dead and H&E staining revealed elongated cells after 14 days of osteogenic induction **(figure 6(d))**, suggesting a temporal regulation of cell attachment in these gels. These dynamic changes may be due to the dissolution of the weak ionic bonds and subsequent increases in mesh size. As expected, collagen content was significantly higher in the alg-gel hydrogels due to the incorporation of gelatin in the bioink. We did not detect differences in collagen deposition among alg-CaCl_2_, alg-CaSO_4_ and alg-ncel bioinks **(figure 6(b**)), perhaps due to the nonfouling nature of the alginate polymer.

**Figure 6.**
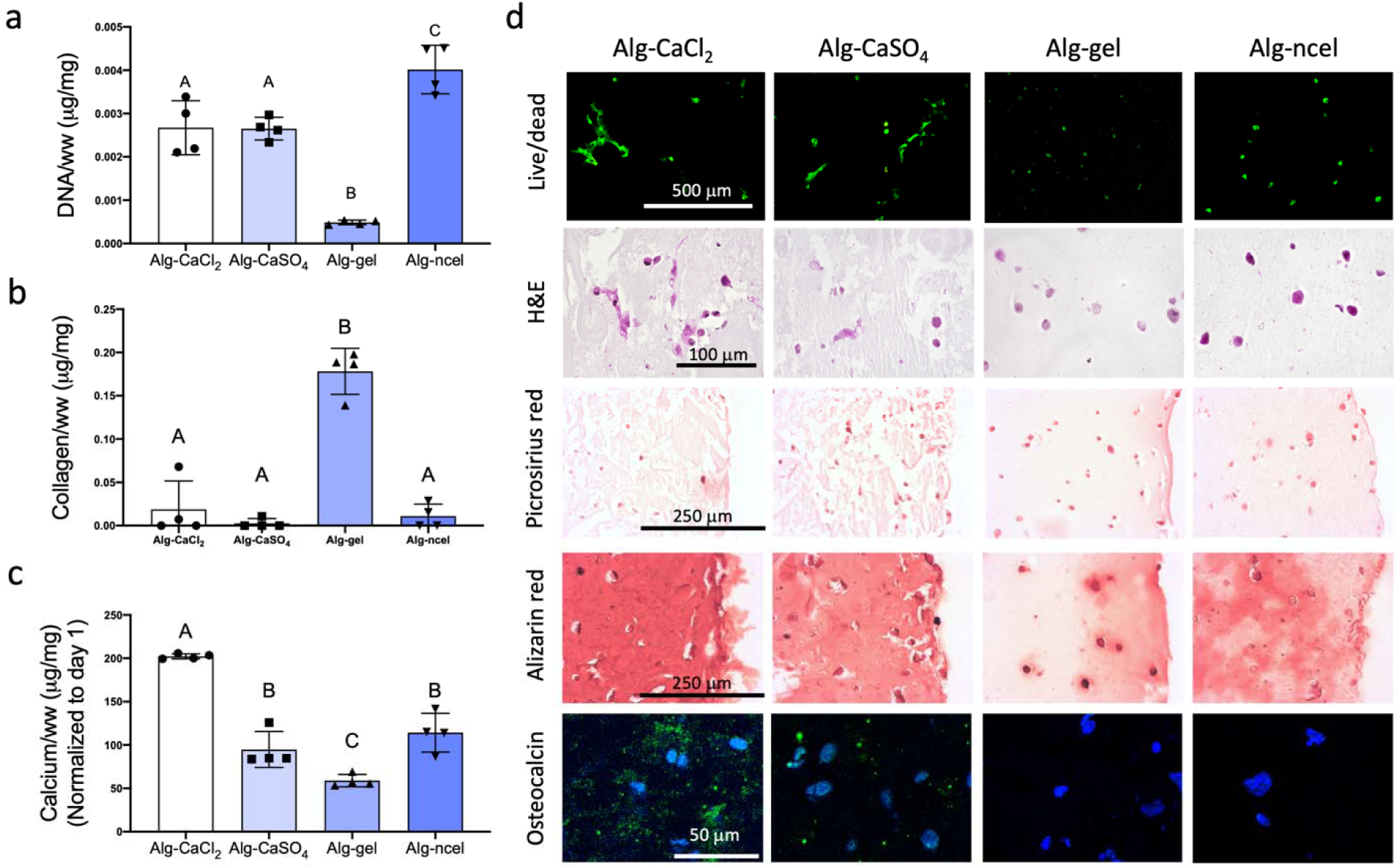
MSC osteogenic potential depends on the type of alginate-based bioink. **(a)** DNA content in 3D printed constructs at day 14 of culture in osteogenic media. **(b)** Collagen content in the 3D printed constructs at day 14. **(c)** Calcium content at day 14 and normalized to day 1 calcium content. **(d)** Live/dead staining and histological examination through staining with H&E, picrosirius red (collagen), Alizarin red (calcium), and IHC for osteocalcin (cell nuclei in blue and osteocalcin in green). n = 4.

### 3.5. Alginate-CaCl_2_ can be used for printing anatomically accurate scaphoid bone substitutes

The scaphoid bone is one of eight independently moving carpal bones in the wrist. Scaphoid fractures account for 60-70% of carpal fractures, with an incidence of 10.6 per 100,000 person-years in the US [50]. Unfortunately, greater than 10% of these fractures progress to non-union [50], which must be treated with bone autografts. Due to the intricate structure and patient-specific shape of the scaphoid bone, 3D printing has been proposed as a clinical tool for the treatment of these fractures and for use in pre-operative planning [51], reconstructive microsurgery [52], and the printing of patient-specific ceramic implants [53]. To 3D print anatomically cell-laden scaphoid grafts, MRI scans from a healthy patient **(figure 7(a))** were used to create a 3D model of the scaphoid shape **(figure 7(b))**, which was sliced **(figure 7(c))** to generate a G-code for printing. Due to improved cell survival, cell-matrix interactions, and MSC osteogenesis identified in this study **(figures 5 and 6)**, we selected the alg-CaCl_2_ bioink to 3D print anatomically accurate cell-laden scaphoid constructs. Once printability was confirmed through 3D printing with Pluronic **(figure 7(d))**, a scaled down (50%) version, which was more convenient for *in vitro* culture, was 3D printed with the alg-CaCl_2_ bioink containing 10×10^6^ MSCs/mL **(figure 7(e))**. Printed constructs were then cultured for 14 days in osteogenic media. MSCs remained viable one day post-printing **(figure 7(f))**. After 14 days in osteogenic media, 3D printed cell-laden scaphoids exhibited a calcified matrix **(figure 7(g))** and positive osteocalcin staining **(figure 7(h))**. Histological examination through H&E of the 3D printed scaphoid cross section revealed cells at the center and periphery of the constructs **(figure 7(i-k))**. Alizarin red staining showed that constructs were not uniformly calcified, with increased calcium deposition observed at the construct periphery **(figure 7(l-n))**. Although the heterogeneous calcification of 3D printed constructs represents a limitation of this current approach, bioreactor culture together with printing of a porous internal architecture to facilitate homogenous nutrient and oxygen diffusion will be explored in future studies.

**Figure 7.**
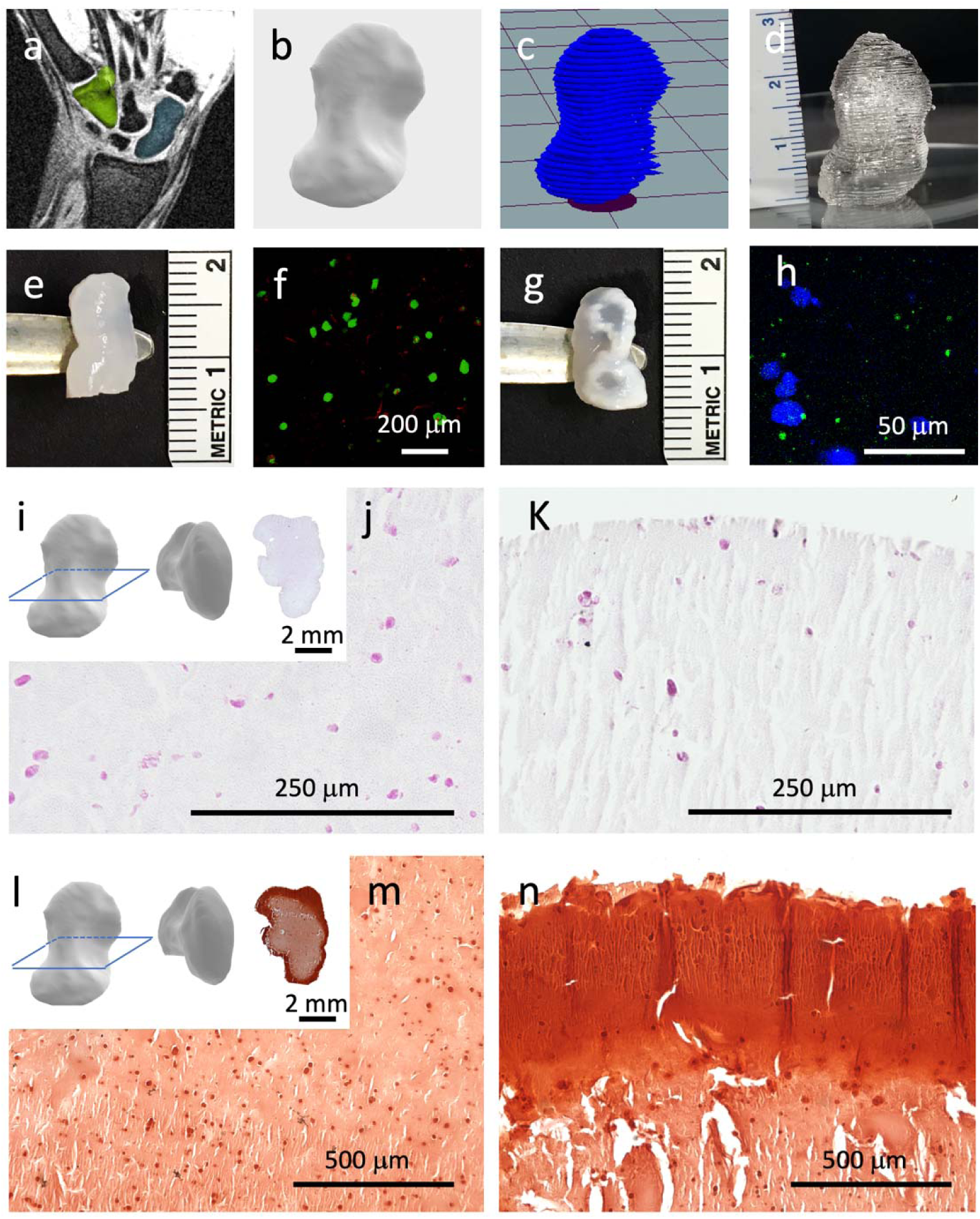
Bioprinting a scaphoid bone replacement. MRI images of the scaphoid of a healthy patient **(a)** (scaphoid in blue, hamate in green) were used to produce a 3D model **(b)**, which was sliced **(c)** for 3D printing. **(d)** 3D printed full-size scaphoid printed with Pluronic. **(e)** Scaled down (50%) alg-CaCl_2_ printed scaphoid after 3D printing and crosslinking. **(f)** Live/dead imaging of MSCs encapsulated in 3D printed scaphoids 1 day post-printing. **(g)** Alg-CaCl_2_ 3D printed scaphoid after 14 days of *in vitro* culture. **(h)** Osteocalcin immunostaining of the 3D printed scaphoids after 14 days of culture (cell nuclei in blue and osteocalcin in green). **(i)** H&E histological analysis of the cross section of osteogenically-induced 3D printed scaphoids at the center **(j)** and periphery **(k)** of the construct. **(l)** Alizarin red staining of 3D printed scaphoids at the center **(m)** and periphery **(n)** of the construct.

The clinical application of 3D printed grafts is limited by poor mechanical properties and lack of initial vascularization. However, these shortcomings could be addressed in a number of ways. For example, mechanical properties of 3D printed tissues could be bolstered by reinforcement through co-printing of biodegradable thermoplastics and cell-laden bioinks [13, 18]. Porosity could be increased by modification of the internal structure of 3D printed scaffolds, which resulted in increased scaffold vascularization after implantation and greater bone regeneration [53, 54]. Moreover, the development of new approaches to dynamically change the mechanical properties post-printing using light or other external stimuli could provide expanded utility for this approach.

## 4. Conclusion

The selection of bioinks is commonly pursued by determination of rheological properties effect on printability. However, the results of this study unequivocally demonstrate that other biophysical properties must be considered when selecting a bioink for 3D printing. Despite similar rheological properties, the four alginate-based bioinks analyzed in this study possessed different physicochemical properties that influenced shape fidelity, construct mechanics and, ultimately, cell function and differentiation. In addition, osteogenic differentiation of entrapped MSCs was dependent upon promoting cell-matrix interactions in 3D printed constructs. 3D bioprinting of a scaphoid bone using the alg-CaCl_2_ bioink also demonstrated the therapeutic potential of this approach to fabricate living osteogenic tissue grafts. These results highlight the importance of assessing not only the rheological properties of bioinks, but also their long-term biological performance for the design of clinically relevant bioinks that can guide cell function towards the desired therapeutic target.

## Disclosure statement

The authors have nothing to disclose.

## Acknowledgements

This work was supported by the National Institutes of Health under award number R01 DE025899 to JKL. The content is solely the responsibility of the authors and does not necessarily represent the official views of the National Institutes of Health. TGF received support from the American Heart Association Postdoctoral Fellowship (19POST34460034). AJT received support from the UC Davis Provost’s Undergraduate Fellowship (PUF) and the California Alliance for Minority Participation (CAMP) Scholarship. This work was also supported by the American Heart Association grant #17IRG33420114 and 19IPLOI34760654 to EAS. The funders had no role in the decision to publish, or preparation of the manuscript. We thank Abhijit Chaudhari for providing the 3D reconstructions of the scaphoid bone.

## Notes

### Competing Interest Statement

The authors have declared no competing interest.

